# Hot Pursuit: Bioinformatic and Biochemical Characterization of a Hyperthermophilic Family B DNA Polymerase from *Pyrolobus fumarii* A1

**DOI:** 10.64898/2026.06.25.734501

**Authors:** Wojciech Rusinek, Sebastian Dorawa, Tadeusz Kaczorowski

## Abstract

Thermostable DNA polymerases are indispensable tools in molecular biology, yet enzymes from the most extreme hyperthermophiles remain largely uncharacterized. Here, we report the biochemical and structural characterization of a family B DNA polymerase from *Pyrolobus fumarii* A1 (Pyrfu pol), one of the most thermoresistant archaea described to date. The enzyme was efficiently overproduced in *E. coli* Rosetta 2(DE3)[pLysS] and purified to homogeneity using a two-step protocol that combined heat treatment with immobilized metal affinity chromatography (IMAC). Bioinformatic analysis confirmed the canonical family B architecture, while AlphaFold-based structural modeling and comparative analysis with mesophilic RB69 DNA polymerase revealed a well-conserved structural core alongside thermoadaptive features. Radiolabel incorporation assays demonstrated enzymatic activity over a broad ionic strength range and an absolute requirement for Mg^2^□. PCR-based optimization confirmed these findings and revealed broad pH tolerance (6.5-11.0). Notably, Tris inhibited radiolabel-based assays (pH 7.0) yet proved essential for efficient PCR amplification (pH 8.5), suggesting a context-dependent role of buffer composition in polymerase activity. Processivity assays confirmed amplification of DNA fragments up to approximately 8,000 bp. Replication fidelity, assessed by the lacZ-based assay, showed a 2.9-fold improvement over Taq polymerase. Urea-nanoDSF yielded an exceptional melting temperature of 105.9 ± 0.08 °C. Pyrfu pol also demonstrated tolerance to common PCR inhibitors, highlighting its potential utility in molecular biology applications.

**Graphical abstract:** 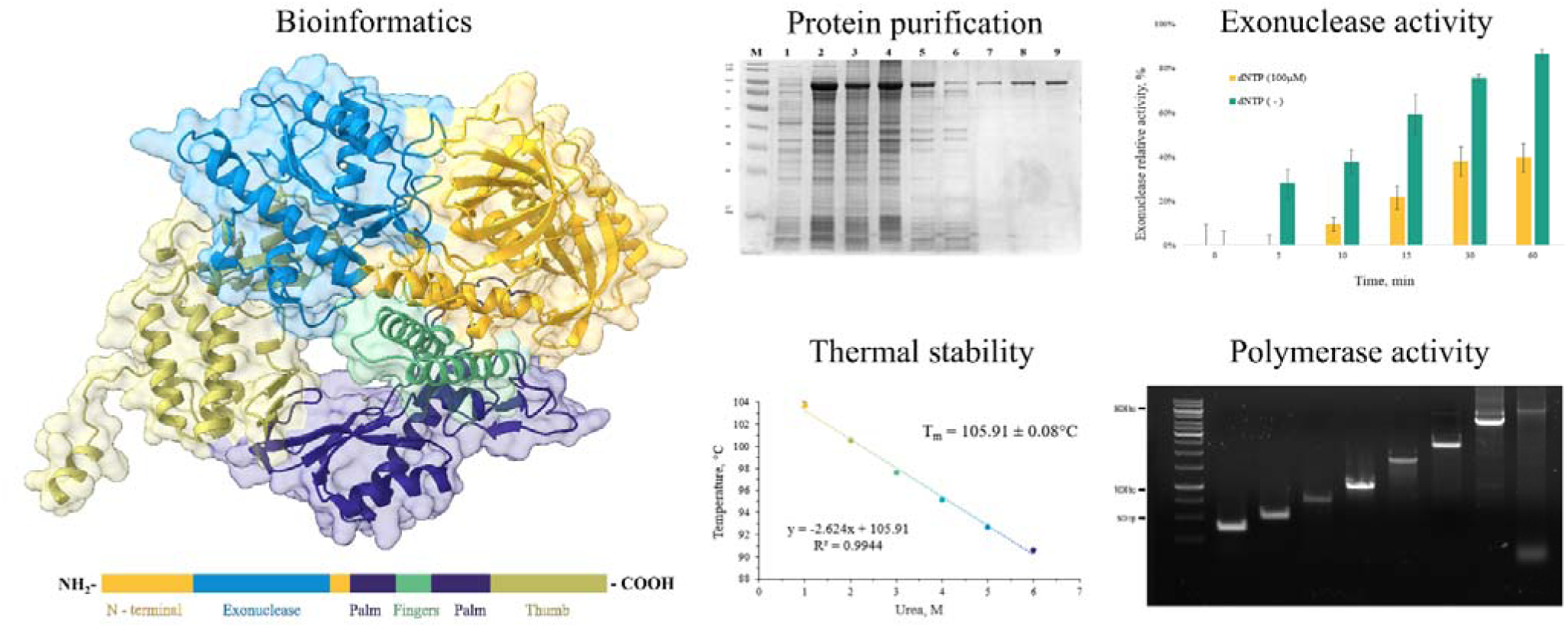

## 1. Introduction

DNA polymerases are highly specialized molecular nanomachines responsible for the faithful replication and maintenance of genetic information, operating with remarkable speed and accuracy [1]. Among them, family B DNA polymerases (PolBs) are among the most widespread, occurring in all three domains of life as well as in numerous DNA viruses [2–3]. In Archaea and Eukarya, they play a central role in genome replication and maintenance, catalyzing highly accurate DNA synthesis and participating in diverse DNA repair pathways [4]. The remarkable fidelity of PolBs results from the coordinated action of two catalytic functions: the 5′→3′ DNA polymerase activity and the intrinsic 3′→5′ exonuclease (proofreading) activity that removes incorrectly inserted nucleotides [5–6].

Despite considerable amino acid sequence diversity among family members, PolBs share a conserved modular architecture composed of three principal domains: an *N*-terminal domain, a 3′→5′ exonuclease (Exo) domain, and a *C*-terminal polymerase (Pol) domain [7]. Conserved catalytic motifs within the Exo (ExoI, ExoII, and ExoIII) and Pol (Motif A, Motif B, Motif C) domains are essential for exonucleolytic proofreading and phosphodiester bond formation, respectively [3,8–9]. While the polymerase and exonuclease functions are relatively well characterized, the role of the *N*-terminal domain remains less understood. In archaeal PolBs, this region has been implicated in uracil sensing and stalling at deaminated bases, a mechanism believed to protect hyperthermophilic genomes from elevated cytosine deamination occurring at high temperatures [10].

Archaea inhabiting extreme geothermal environments possess replication systems adapted to function under conditions that challenge the stability of nucleic acids and proteins. Consequently, archaeal DNA polymerases, particularly those from hyperthermophiles, have attracted substantial attention for their extraordinary thermostability, processivity, and fidelity [11]. Several archaeal family B polymerases, including enzymes from the genera *Pyrococcus* and *Thermococcus*, have become indispensable tools in molecular biology and biotechnology, especially for high-fidelity PCR applications [12–14]. Nevertheless, despite extensive studies of model archaeal polymerases, the diversity and biochemical properties of PolBs from many extremophilic archaeal lineages remain poorly characterized.

One particularly intriguing lineage of hyperthermophilic archaea is the genus *Pyrolobus*, whose representatives inhabit deep-sea hydrothermal environments characterized by extreme temperatures, hydrostatic pressure, and geochemical conditions [15]. Among them, *Pyrolobus fumarii*, an obligate anaerobe originally isolated from deep-sea hydrothermal vents (black smokers) on the Mid-Atlantic Ridge, is recognized as one of the most thermophilic organisms known to date, exhibiting growth up to 113 °C and an optimal growth temperature near 106 °C [16]. The extreme environmental niche occupied by *P. fumarii* suggests that its replication machinery possesses unique molecular adaptations that enable efficient metabolism under conditions of over-boiling. Such adaptations may provide valuable insights into the evolution of thermostable enzymes and offer promising candidates for biotechnological applications requiring robust DNA-processing activities.

In this study, we present a bioinformatic and biochemical characterization of a family B DNA polymerase from *P. fumarii* strain A1. The analyzed enzymes display the conserved structural hallmarks of archaeal PolBs and may harbor adaptations associated with a hyperthermophilic lifestyle. By combining sequence analysis, structural predictions, and biochemical assays, we aimed to investigate its catalytic properties, thermostability, and functional features relevant to DNA replication fidelity and potential biotechnological utility.

### 2. Materials and methods

#### 2.1 Bioinformatic analysis

The amino acid sequence of Pyrfu pol was retrieved from the NCBI protein database (accession no. WP_014025844.1) [17]. Physicochemical parameters of the native protein (803 aa) and the His-tagged variant (818 aa) were calculated using the ProtParam tool (ExPASy) [18]. Domain architecture was determined using InterPro (EMBL-EBI) [19]. Amino acid composition was compared between Pyrfu pol and the mesophilic RB69 phage DNA polymerase to investigate compositional differences potentially associated with the thermostability of Pyrfu pol. The three-dimensional structure of Pyrfu pol (803 aa) was predicted using the AlphaFold 3 server [20]. Modeling accuracy was evaluated using the pTM parameter. Protein visualization and domain coloring were performed in UCSF ChimeraX v1.10 [21]: the *N*-terminal domain was colored gold, the 3′→5′exonuclease domain light blue, the palm domain navy blue, the fingers domain fern green, and the thumb domain olive. Multiple sequence alignment was generated using Clustal Omega v1.2.4 (EMBL-EBI), and sequence similarity was visualized using Bioedit v7.2 [22].

### 3.2 Structural comparison with (mesophilic) RB69 phage DNA polymerase

The predicted structure of Pyrfu pol was superimposed onto the crystal structure of the RB69 phage DNA polymerase (PDB: 1WAF, 903 aa) using the matchmaker command in UCSF ChimeraX v1.10. Structural alignment was performed with the Needleman-Wunsch algorithm using the BLOSUM-62 substitution matrix, an SS fraction of 0.3, and gap open penalties of 18/18/6 (helix/strand/other) with a gap extension penalty of 1. Per-residue Cα RMSD values were calculated for all aligned residue pairs and mapped onto the Pyrfu pol backbone as a color gradient ranging from golden (RMSD ≈ 0 Å) to purple (RMSD ≈ 29 Å). Structural elements unique to one of the two polymerases, with no equivalent counterpart in the aligned structure, were depicted in gray.

The secondary-structure overview of both polymerases was determined using ChimeraX. Residues were classified into three categories: alpha-helices, beta-strands, and coils/turns. Quantification was performed by isolating specific structural types using the commands: select helix, select strand, and select coil, and by calculating their occupancy relative to the total number of residues in each model (803 aa for Pyrfu pol and 903 aa for RB69 pol).

### 2.3 Bacterial strain, media, culture conditions, primers and plasmids

The *E. coli* RosettaTM 2(DE3)[pLysS] (Novagen, Germany) was used for protein overproduction. Cells were grown in Lysogenic broth (LB) or on Lysogenic agar (LA) plates supplemented with kanamycin (50 µg/mL). Protein overproduction was carried out using the expression vector pET24a(+)_Pyrfu (BioCat, Germany). Oligonucleotides were designed using SnapGene software (www.snapgene.com) and refined using PCR Primer Stats [23] based on the pET24a(+)_Pyrfu (7,664 bp) and pET24a(+)_Pya (7,719 bp) constructs (**Table 1**).

**Table 1.**
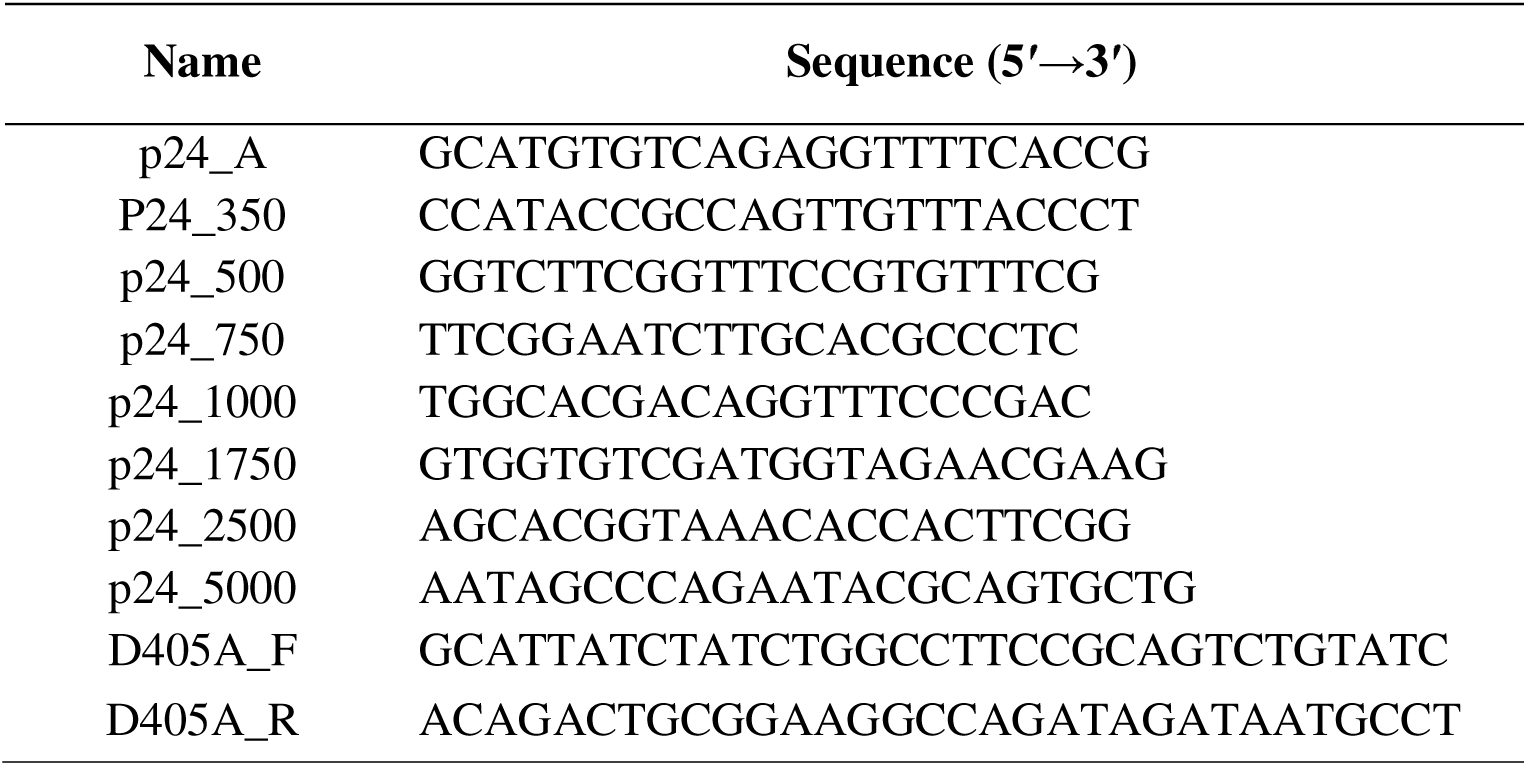
Sequences of oligonucleotides used in this work.

### 2.4 Overproduction and purification of the recombinant protein

Pyrfu DNA polymerase was overproduced in *E. coli* Rosetta 2(DE3)[pLysS] harboring the expression plasmid. The culture was grown in 300 mL of LB medium supplemented with kanamycin at 37 °C to an OD600 of 0.7. Protein overproduction was induced by adding IPTG to a final concentration of 0.5 mM (Serva, Germany), followed by incubation for 2 h at 37 °C. Cells were harvested by centrifugation (4,000 × g, 20 min, 4 °C) and stored at -80 °C.

The cell pellet was resuspended in buffer A (50 mM NaH_2_PO_4_ pH 8.0, 300 mM NaCl, 0.1% [v/v] Triton X-100, 5% [v/v] glycerol, and 10 mM imidazole) and disrupted by sonication on ice (42 cycles of 10 s at an amplitude of 69%) using a Branson SFX550 sonicator (Emerson Electric Corp, USA). Cell debris was removed by centrifugation (10,000 × g, 20 min, 4 °C). The supernatant was incubated at 90 °C for 20 min to denature thermolabile host proteins, then centrifuged again (10,000 × g, 20 min, 4 °C) to obtain a clarified lysate. The clarified lysate was applied to a 1.5 mL TALON Metal Affinity Resin (Takara Bio, USA). The resin was washed with 15 mL of buffer A, followed by an additional 15 mL of buffer B (buffer A supplemented with 20 mM imidazole). Bound proteins were eluted with buffer C (buffer A supplemented with 150 mM imidazole). Fractions containing the target protein were pooled and dialyzed against storage buffer (10 mM Tris (pH 8.0), 50 mM KCl, 1 mM DTT, 0.1 mM EDTA, 0.5% Triton X-100, and 50% [v/v] glycerol), then stored at -20 °C for further analysis. All purification steps were monitored by SDS-PAGE, and protein concentrations were determined using the Bradford assay [24].

### 2.5 Pyrfu pol activity assay using radiolabeled nucleotides

Pyrfu activity was assayed by measuring incorporation of radiolabeled nucleotides into calf-thymus DNA. The reaction mixture (50 µL) consisted of 5 µL of buffer D (20 mM KCl, 2 mM MgSO_4_, and 20 mM (NH_4_)_2_SO_4_ with the concentrations of Tris (pH 7.0), KCl, MgSO_4_, and (NH_4_)_2_SO_4_ varied independently across a gradient, alongside reaction pH (6.0-10.0) and incubation temperature (55-90 °C). The buffer was supplemented with 200 µM dATP, 200 µM dCTP, 200 µM dGTP, 20 µM dTTP, 3 µCi/mL of [methyl-³H]-thymidine 5’-triphosphate (84.2 Ci/mmol; PerkinElmer), 1µL of Pyrfu pol (1 mg/mL), and 0.2 mg/mL of activated calf thymus DNA. The reactions were initiated by adding aliquots of Pyrfu pol on ice; samples were then immediately transferred to a preheated thermal block and incubated for 10 min at 80 °C, after which the reaction was stopped by adding 200 µL of 10% [w/v] trichloroacetic acid (TCA) and kept on ice for 10 min. After termination, an aliquot was spotted on a GF/C filter disc (Whatman, Clifton, USA). The filters were washed three times with 5 mL of 5% [w/v] TCA, then twice with 5 mL of 70% [v/v] ethanol. The filters were air-dried and counted in a liquid scintillation cocktail (MP Biomedicals, USA) using a scintillation counter (PerkinElmer, USA). All measurements were performed in triplicate.

### 2.6 Pyrfu pol 3′→5′ exonuclease activity assay using radiolabeled nucleotides

The exonuclease activity of Pyrfu pol was assessed by monitoring degradation of a radiolabeled substrate. Briefly, Lambda phage DNA digested with HindIII (Thermo Fisher Scientific, USA) was labeled with [³H]-dTTP using the Klenow fragment. The reaction mixture (50 µL) was prepared as described for the polymerase activity assay, except that the Pyrfu pol concentration was 0.5 mg/mL. Reactions were carried out under optimized buffer and temperature conditions, with or without dNTPs, and were terminated at 0, 5, 10, 15, 30, and 60 min by adding 200 µL of 10% [w/v] TCA. Exonuclease activity was expressed as the decrease in TCA-insoluble radioactivity over time, reflecting the progressive release of labeled nucleotides resulting from DNA degradation. TCA precipitation, filtration, and scintillation counting were performed as described above. All measurements were performed in triplicate

### 2.7 Pyrfu pol activity assay using Polymerase Chain Reaction (PCR)

Pyrfu pol activity was analyzed by PCR and evaluated by agarose gel electrophoresis. Reaction conditions were optimized by independently varying reaction pH (6.0-11.0), MgSO_4_ (0-10 mM), KCl (0-100 mM), (NH_4_)_2_SO_4_ (0-100 mM), Tris (pH 8.5, 0-100 mM), and extension rate. The condition that produced the clearest band on agarose gel was selected as optimal. The final buffer composition (1X) was as follows: 40 mM Tris (pH 8.5), 20 mM KCl, 2 mM MgSO_4_, 20 mM (NH_4_)_2_SO_4_, and 0.1% Triton X-100. Extension rates were optimized based on product size: 20 s/kb for products up to 500 bp, 40 s/kb for products between 500 bp and 5,000 bp, and 60 s/kb for products exceeding 5,000 bp.

The 50 µL reaction mixture contained 5 µL of reaction buffer (×10), 1 µL of 5 mM dNTPs (Thermo Fisher Scientific, USA), 1 µL of 10 µM forward primer (p24_A, D405A_F, Table 1), 1 µL of 10 µM reverse primer (p24_500, p24_1000, D405A_R, Table 1) (Genomed, Poland), and 8 ng of plasmid DNA pET24a(+)_Pyrfu as template for products up to 5,000 bp and pET24a(+)_Pya as template for ∼8,000 bp product. PCR was performed using the following thermal cycling profile: an initial denaturation at 98 °C for 2 min, followed by 30 cycles of denaturation at 98 °C for 15 s, annealing at a primer-specific temperature for 20 s, and extension at 72 °C for a time adjusted to product size. PCR products were separated by electrophoresis on a 0.8% agarose gel in TBE buffer and visualized by ethidium bromide staining; a 1 kb GeneRuler (Thermo Fisher Scientific, USA) was used as a size reference.

### 2.8 Analysis of Pyrfu pol processivity using PCR amplification

The processivity of Pyrfu pol was evaluated by PCR amplification of plasmid DNA fragments of increasing length: 350, 500, 750, 1,000, 1,750, 2,500, 5,000, and ∼8,000 bp. Reaction mixtures contained 1 µL of 10 µM forward primer (p24_A, D405A_F; Table 1), 1 µL of 10 µM reverse primer (p24_350, p24_500, p24_750, p24_1000, p24_1750, p24_2500, p24_5000, and D405A_R; Table 1), and 8 ng of plasmid DNA pET24a(+)_Pyrfu was used for products 350-5,000 bp, and pET24a(+)_Pya was used for the ∼8,000 bp product. All reactions were performed under optimized conditions as described above, with extension time adjusted according to product size. Processivity was defined as the maximum fragment length yielding a detectable PCR product, as determined by agarose gel electrophoresis. PCR products were separated on a 0.8% agarose gel in TBE buffer and visualized by ethidium bromide staining; a 1 kb GeneRuler molecular weight marker (Thermo Fisher Scientific) was used as a size reference.

### 2.9 Resistance of Pyrfu pol to PCR-inhibiting agents

The resistance of Pyrfu pol to common PCR inhibitors was assessed by agarose gel electrophoresis. Each inhibitor was added independently to the reaction mixture at the following concentration: NaCl (0-60 mM), heparin (0-75 ng per reaction), lactoferrin (0-30 µg per reaction), humic acid (0-75 ng per reaction), EDTA (0-5 mM), phenol (0-4% [v/v]), ethanol (0-5% [v/v]), and isopropanol (0-5% [v/v]). The reaction mixture (50 µL) contained 1 µL of 5 mM dNTPs, 1 µL of 10 µM forward primer p24_A (Table 1), 1 µL of 10 µM reverse primer p24_500 (Table 1), and 8 ng of pET24a(+)_Pyrfu plasmid DNA as template in the optimized buffer (40 mM Tris (pH 8.0), 20 mM KCl, 2 mM MgSO_4_, 20 mM (NH_4_)_2_SO_4_, and 1% Triton X-100). The amplification target was a 500 bp product. PCR was performed using the following thermal cycling profile: initial denaturation at 98 °C for 2 min, followed by 32 cycles of denaturation at 98 °C for 15 s, annealing at a primer-specific temperature for 20 s, and extension at 72 °C for 10 s. PCR products were resolved on a 0.8% agarose gel and visualized by ethidium bromide staining; a 1 kb GeneRuler served as a size reference.

### 2.10 Pyrfu pol fidelity assay using blue/white colony screening

Fidelity of DNA polymerase was evaluated using a plasmid-based gap-filling assay, adapted from Keith et al. [25] with minor adjustments. The 20 µL reaction mixture contained 2 mM MgSO_4_, 20 mM KCl, 20 mM (NH_4_)_2_SO_4_, and 250 µM of each dNTP. Final concentrations were 100 nM for Pyrfu pol and 10 nM for gapped pSJ3 plasmid DNA. Gap-filling reactions proceeded at 80 °C for 10 min. Products were then used directly to transform chemically supercompetent *E. coli* DH5α cells. Transformants were plated on LA medium containing ampicillin (100 μg/mL), X-gal (40 μg/mL), and IPTG (1 mM), then incubated at 37 °C for 16 hrs. Mutation frequency was determined by blue/white colony screening. Background mutation rates were derived from no-polymerase control reactions. In addition, reactions performed with Taq DNA polymerase and Pfu DNA polymerase served as low- and high-fidelity reference controls, respectively. Mutation frequencies and error rates were calculated as per the referenced method.

### 2.11 Melting temperature analysis of Pyrfu pol by urea-nanoDSF

Thermal stability was assessed by urea-nanoDSF (urea-gradient nano differential scanning fluorimetry) following the procedure described by Rusinek and Dorawa [26]. Measurements were performed on a Prometheus NT.48 instrument (NanoTemper Technologies, Germany). Standard-grade NT.48 glass capillaries were filled with 10 μL of recombinant protein stock solution, supplemented with 90 μL of 1-6 M urea. Prior to analysis, samples were equilibrated for 5 min at room temperature, then centrifuged at 4,000 × g for 5 min. Samples were then heated from 20 °C to 110 °C at a rate of 0.5 °C/min. Protein unfolding was monitored by intrinsic tryptophan and tyrosine fluorescence at emission wavelengths of 350 and 330 nm, respectively. Data were analyzed using PR.StabilityAnalysis software. Melting temperatures were determined as the maxima of the first derivative of the fluorescence ratio (F_350_/F_330_), and the final thermal stability in the absence of denaturant was calculated by linear extrapolation of the T_m_ values obtained across the urea concentration gradient (1-6 M). All measurements were performed in triplicate.

## 3. Results and Discussion

### 3.1 Bioinformatic analysis

Computational analysis of Pyrfu pol using the ProtParam tool revealed that the native protein (803 aa) has a theoretical molecular weight of 92,68 kDa and an isoelectric point of 8.85, classifying it as a moderately basic protein. The instability index of 38.73 indicates that the protein is stable. The aliphatic index of 95.48, reflecting the relative volume occupied by aliphatic side chains, suggests a high degree of hydrophobic core packing, which could contribute to protein thermostability. The negative GRAVY value (−0.300) indicates an overall hydrophilic character of the protein, suggesting that it is soluble and functional in aqueous environments. The His-tagged variant (818 aa) displayed nearly identical parameters (MW 94,41 kDa, pI 8.80, instability index 37.62, aliphatic index 95.40, GRAVY −0.310), suggesting that the addition of the affinity tag does not significantly alter the overall physicochemical profile of the protein. Domain analysis using InterPro confirmed that Pyrfu pol belongs to the family B DNA polymerases (IPR050240, IPR006172), consistent with its origin and the known distribution of family B polymerases among Archaea. Two major functional domains were identified: the family B DNA polymerase (nucleotidyltransferase) domain (PF00136, IPR006134, residues 406-803), responsible for template-directed DNA synthesis, and the 3’→5’ exonuclease domain (PF03104, IPR006133, residues 178-365), which provides proofreading activity. Within the polymerase domain, three structural subdomains were found: the palm subdomain (IPR023211, residues 406-485 and 537-623), the fingers subdomain (G3DSA:1.10.287.690, residues 486-536), and the thumb subdomain (IPR042087, residues 624-803) as well as N-terminal domain (domain 1, G3DSA:3.30.342.10 residues 1-177, 366-405), reflecting the canonical “right-hand” architecture characteristic of all family B DNA polymerases [27–28]. The identification of conserved catalytic motifs ExoI, ExoII, and ExoIII within the exonuclease domain, and motifs A, B, and C within the palm and fingers subdomain (**Figure 1c**), was confirmed by the PROSITE signature PS00116. Gene Ontology annotation assigned Pyrfu pol to DNA-dependent DNA polymerase activity (GO:0003887) and DNA replication (GO:0006261), consistent with the enzymatic activity demonstrated experimentally in this study.

**Figure 1.**
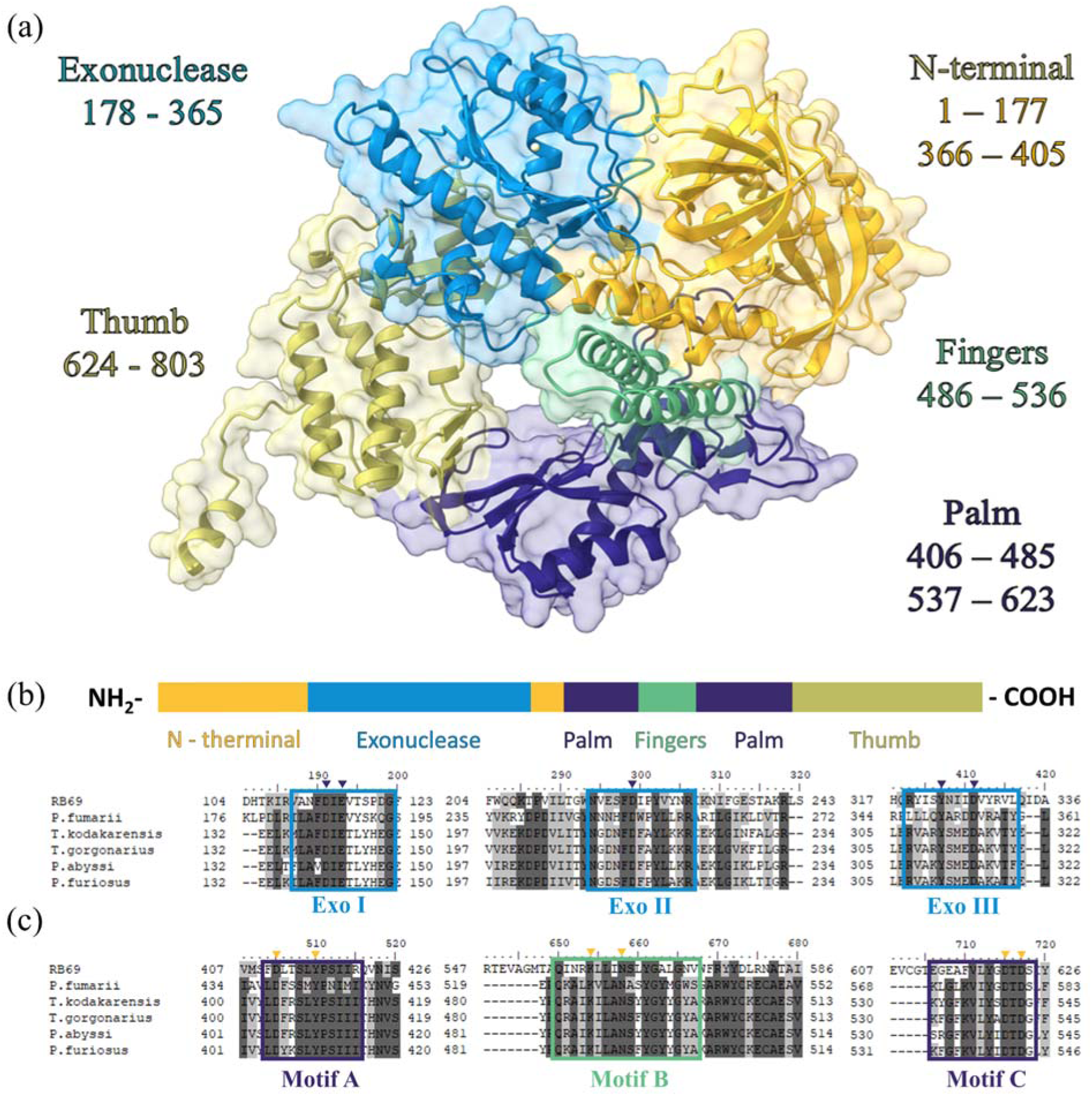
Predicted three-dimensional structure and domain architecture of Pyrfu pol. **(a)** The structural model was generated using AlphaFold 3 (pTM = 0.91) and visualized in UCSF ChimeraX. Domains were color-coded and mapped based on InterPro architecture analysis, as follows: N-terminal domain (gold), exonuclease domain (light blue), palm domain (navy blue), fingers domain (fern green), and thumb domain (olive). **(b)** Linear domain map of Pyrfu pol showing the arrangement of domains along the polypeptide chain (NH to COOH): N-terminal domain (residues 1-177 and 366-405), exonuclease domain (residues 178-365), palm subdomain (residues 406-485 and 537-623), fingers subdomain (residues 486-536), and thumb subdomain (residues 624-803). **(c)** Multiple sequence alignment of Pyrfu pol with representative family B DNA polymerases (RB69 gp43, *P. fumarii*, *T. kodakarensis*, *T. gorgonarius*, *P. abyssi, P. furiosus*), highlighting conserved sequence motifs of the exonuclease domain (Exo I, Exo II, Exo III) and the polymerase domain (Motif A, Motif B, Motif C). Catalytic residues are indicated by triangles above the alignment. Light gray indicates 70% sequence identity and dark gray indicates 100% sequence identity.

The three-dimensional structure of Pyrfu pol was predicted using the AlphaFold 3 server, which represents the current state-of-the-art in de novo protein structure prediction (**Figure 1a**). The model achieved a global pTM score of 0.91, indicating high confidence in the prediction. Per-residue model quality, assessed by the pLDDT score (*data not shown*), confirmed that the vast majority of residues received scores above 70, falling within the “Confident” and “Very high” categories. Regions of lower confidence, reflected by yellow and orange coloring in the pLDDT map, were limited to short internal loops in the N-terminal and Fingers subdomains as well as the His-tag sequence, which is expected given the inherent flexibility of loop regions and the absence of structural constraints on affinity tags. The Expected Position Error map provided by the server (*data not shown*) further supported the model’s internal consistency, showing consistently low inter-residue positioning error across the entire protein without distinct blocks of elevated uncertainty, indicating that the predicted relative domain arrangements are reliable. The quality metrics suggest that the AlphaFold 3 model of Pyrfu pol provides a sufficiently accurate structural representation to support the comparative and functional analyses performed in this study.

### 3.3 Structural comparison with (mesophilic) RB69 phage DNA polymerase

A comparison of the amino acid composition of Pyrfu pol with that of the PolB homolog of the mesophilic phage RB69 (RB69 gp43) revealed several differences that may be relevant to its thermoadaptation. Pyrfu pol shares 18.26% sequence identity with RB69 pol and is enriched in arginine (8.3% vs. 5.1%) and depleted in serine (3.5% vs. 6.1%) relative to RB69 gp43. The arginine-for-serine substitution pattern is a hallmark of thermophilic proteins, as arginine forms stronger and more extensive hydrogen-bond networks and salt bridges than serine, thereby contributing to increased structural rigidity at elevated temperatures [29]. Furthermore, Pyrfu pol contains more proline residues (5.9% vs. 4.8%), consistent with the known role of proline in increasing overall protein rigidity, restricting backbone conformational freedom, and decreasing the entropy of unfolding [11].

The structural superimposition of Pyrfu pol onto RB69 DNA polymerase (PDB: 1WAF, **Figure 2 top panel**) was performed in UCSF Chimera X v. 1.10, yielding an overall Cα RMSD of 8.700 Å across all 705 aligned residue pairs, which decreased to 1.240 Å when calculated over 132 pruned, well-aligned pairs. This indicates that despite their evolutionary distance and the markedly different living conditions of their host microbes, the two enzymes share a well-conserved structural core. The per-residue RMSD analysis (**Figure 2 bottom panel**) revealed a pattern of structural conservation that reflects the functional importance of specific regions. The lowest RMSD values were observed within the conserved catalytic motifs. In the exonuclease domain, motifs ExoI, ExoII, and ExoIII displayed the smallest deviations not only relative to each other but also compared to the surrounding inter-motif regions of the exonuclease domain, underscoring the strong evolutionary pressure to maintain the geometry of the proofreading active site. Similarly, polymerase motifs A and C within the palm and motif B in the fingers subdomain exhibited the lowest structural deviation, consistent with their essential role, which is conserved across family B DNA polymerases [30]. In contrast, the highest per-residue Cα RMSD values were observed in inter-motif loop regions distributed throughout all polymerase (sub)domains. Notably, although isolated peaks of high RMSD were scattered across all polymerase (sub)domains, the thumb subdomain displayed a distinct pattern of uniformly elevated deviation across extended consecutive stretches, suggesting a more globally divergent fold in this region. Unique structural elements, absent in the aligned counterpart and shown in gray, were most prominent in the *N*-terminal, palm, fingers, and exonuclease (sub)domains, whereas the thumb domain contained comparatively fewer such elements. These structurally divergent regions likely reflect Pyrfu pol adaptations to the extreme living conditions of *Pyrolobus fumarii*, which grows at temperatures up to 113 °C [16], in stark contrast to the mesophilic *E. coli* bacteriophage RB69. The concentration of structural divergence in non-catalytic, peripheral regions, while catalytic motifs remain conserved, is consistent with a pattern in which functional constraints preserve the active-site architecture across distant homologs, while structural plasticity in flanking regions allows more accommodating organism-specific (e.g., thermostability) adaptations to extreme environments. Notably, a comparison of secondary structure composition in ChimeraX revealed that Pyrfu pol contains a higher proportion of β-strands than RB69 pol (20.2% vs. 11.3%, respectively), despite highly similar α-helical content (40.2% vs. 40.9%). An increased β-strand content has been previously associated with thermostability in family B DNA polymerases, as observed in the thermophilic *Thermococcus gorgonarius* polymerase compared to the mesophilic RB69 phage polymerase [31], suggesting that an analogous structural strategy may also contribute to the exceptional thermal stability of Pyrfu pol (T_m_ = 105.9 ± 0.08 °C).

**Figure 2.**
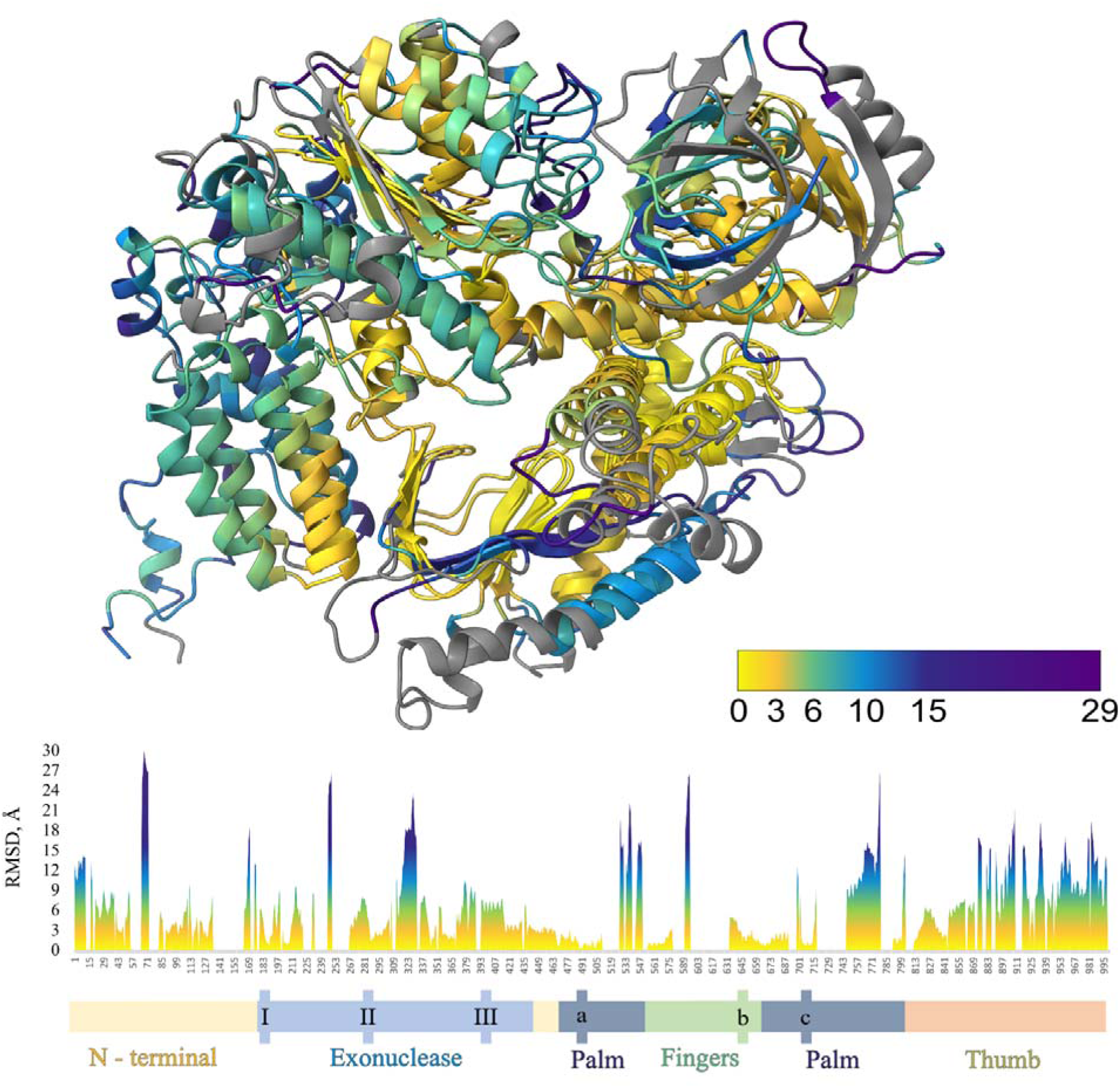
Structural superimposition of Pyrfu pol and RB69 DNA polymerase. The predicted structure of Pyrfu pol (AlphaFold 3) was superimposed onto the crystal structure of RB69 DNA polymerase (PDB: 1WAF) using UCSF ChimeraX (MatchMaker, Needleman-Wunsch algorithm, BLOSUM-62 matrix). The Cα RMSD is mapped per residue onto the Pyrfu pol backbone and color-coded on a gradient from yellow (RMSD ≈ 0 Å) to purple (RMSD ≈ 29 Å). Gray regions indicate structural elements unique to one of the two polymerases, with no equivalent counterpart in the aligned structure. The lower panel shows the per-residue Cα RMSD plotted along the sequence of Pyrfu pol, with the domain map indicated at the bottom. Within the exonuclease domain, conserved motifs ExoI, ExoII, and ExoIII are indicated (I-III); within the palm and finger subdomain, conserved polymerase motifs A, B, and C are indicated (a-c).

### 3.4 Overproduction and Purification of the recombinant protein

Pyrfu pol was efficiently overexpressed in *E. coli* Rosetta 2 (DE3)[pLysS] and successfully purified using Talon Chromatography Resin. A two-step procedure was used to purify the recombinant Pyrfu pol with a His-tag at the *C*-terminus. It consisted of heat treatment (90 °C, 20 min) followed by metal affinity chromatography. At each step, the purity of the enzyme preparation was monitored by 10% SDS-PAGE (**Figure 3**), which revealed a single band of the expected molecular weight. The final preparation yielded approximately 9 mg of purified protein from 300 mL of bacterial culture, indicating an efficient purification procedure suitable for subsequent biochemical characterization.

**Figure 3.**
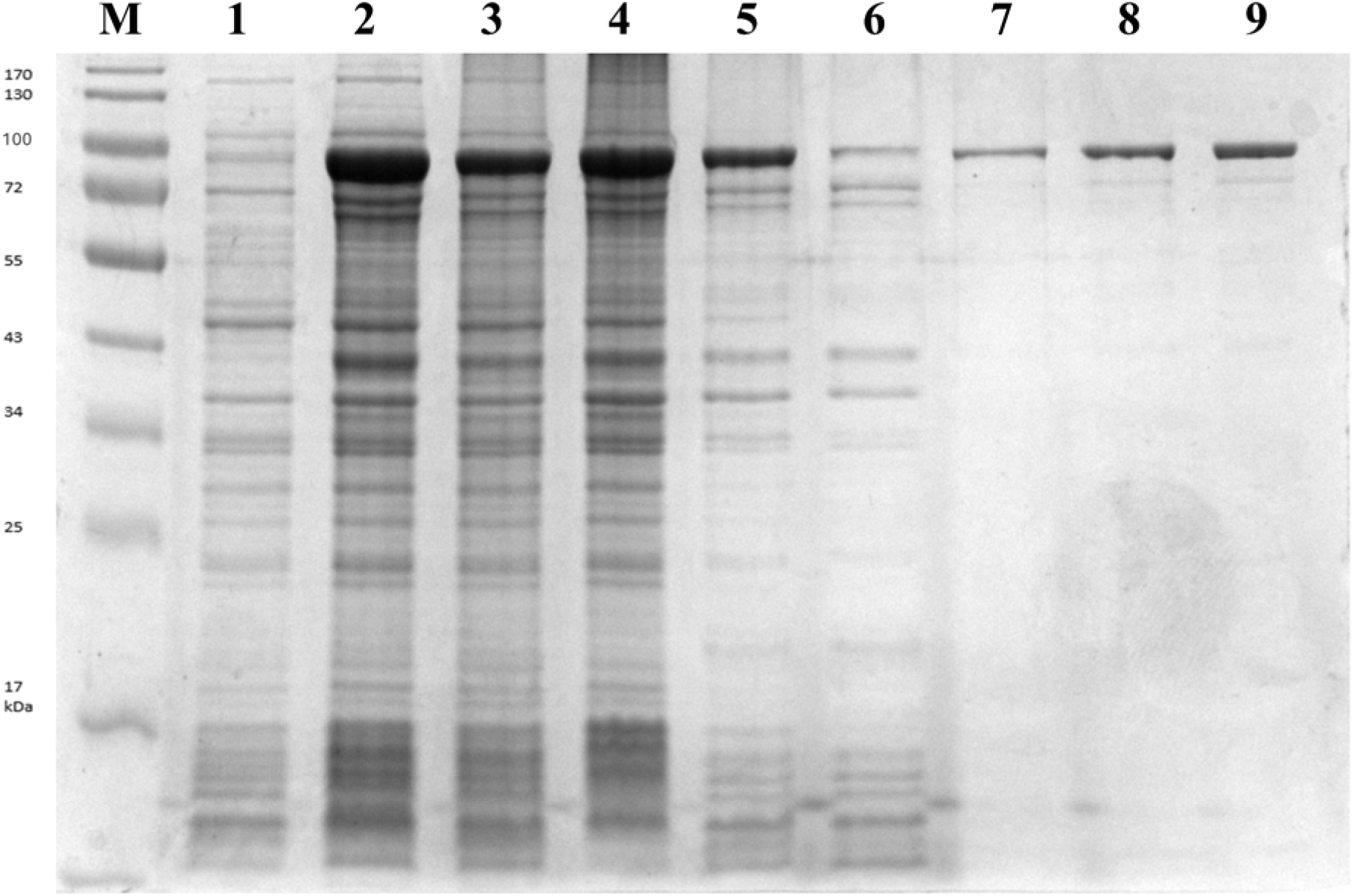
SDS-PAGE analysis of overproduction and purification of Pyrfu pol. Molecular masses (kDa) of marker proteins are indicated on the left side of the gel; the expected molecular weight of the recombinant Pyrfu pol is approximately 94.4 kDa. M: PageRuler Prestained Protein Ladder (Thermo Fisher Scientific); lane 1: whole cell lysate before IPTG induction; lane 2: whole cell lysate after IPTG induction; lane 3: crude lysate; lane 4: pellet fraction after heat treatment; lane 5: supernatant after heat treatment; lane 6: flow-through fraction; lane 7: wash fraction; lane 8: elution fraction; lane 9: protein after dialysis.

### 3.5 Pyrfu pol activity assay using radiolabeled nucleotides

Recombinant enzyme activity was measured by quantifying the incorporation of [³H]-dTTP into activated calf thymus DNA under various experimental conditions. The effect of temperature was tested over 55-90 °C, and the polymerase was active across this range, with highest activity at 80 °C (**Figure 4a**). This is consistent with the hyperthermophilic nature of *Pyrolobus fumarii*, which grows optimally at 106 °C, suggesting that the polymerase retains maximal catalytic efficiency at temperatures well below the organism’s optimal growth temperature, as has been observed for other archaeal family B polymerases [32–33]. The effect of pH was tested in the range of 6.0-10.0 (**Figure 4b**). The highest activity was observed between pH 7.0 and 8.0, with sharp declines at pH values below 6.5 and above 8.5, indicating a relatively narrow pH optimum typical of archaeal replicative polymerases [34]. The requirement for divalent cations was assessed by varying MgSO_4_ concentration from 0 to 10 mM (**Figure 4c**). No activity was detected in the absence of Mg²[, confirming its essential role as a cofactor for catalysis. Activity increased steeply up to 2 mM, then declined gradually at higher concentrations. The optimal MgSO_4_ concentration of 2 mM falls within the range reported for Pfu and Tli thermophilic family B polymerases [14, 35]. The effect of monovalent salts was tested by independently varying KCl (0-100 mM) and (NH_4_)_2_SO_4_ (0-100 mM) concentrations (**Figure 4d-e**). Pyrfu pol displayed remarkably broad tolerance to both salts, retaining over 80% relative activity across the entire tested range for KCl, and showing only a gradual decline in activity at (NH_4_)_2_SO_4_ concentrations above 20 mM, with approximately 50% activity retained at 100 mM.

**Figure 4.**
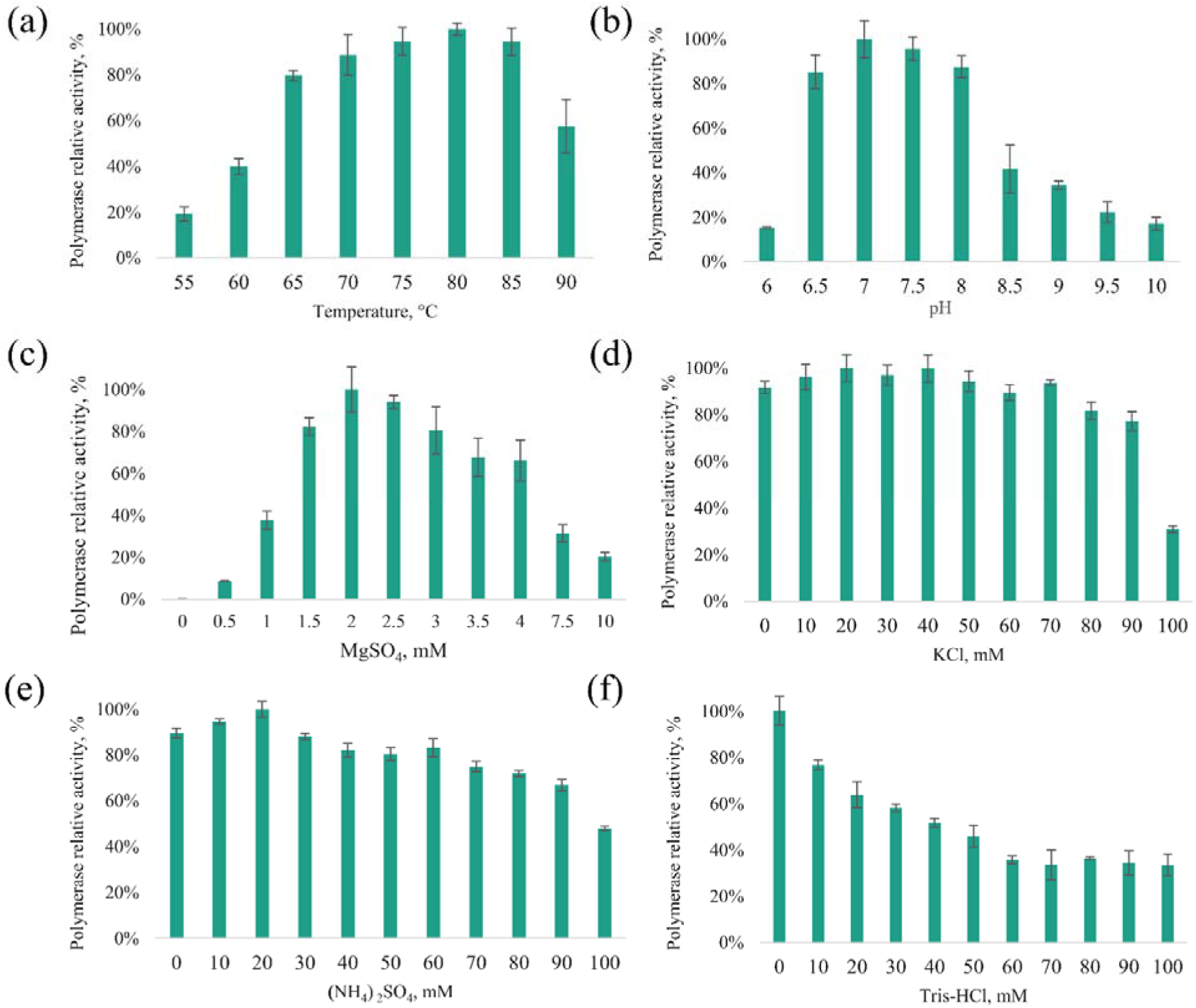
Optimization of reaction conditions for Pyrfu pol. Polymerase activity was measured by incorporation of [³H]-dTTP and expressed as the amount of radioactivity incorporated per reaction. Each parameter was varied independently: (a) reaction temperature (55-90 °C), (b) pH (6.0-10.0), (c) MgSO_4_ (0-10 mM), (d) KCl (0-100 mM), (e) (NH_4_)_2_SO_4_ (0-100 mM), (f) Tris (pH 7.0, 0-100 mM).

The effect of Tris (pH 7.0) was tested over 0-100 mM (**Figure 4f**). A pronounced inhibitory effect was observed in the radiolabel incorporation assay, with activity dropping to approximately 50% at 40 mM and declining further at higher concentrations; the highest activity was observed in the absence of Tris. This observation is noteworthy, as Tris is routinely used in polymerase reaction buffers; however, PCR-based optimization revealed a contrasting requirement, in which a minimum of 10 mM Tris (pH 8.5) was necessary to obtain a barely detectable amplification product, with optimal PCR performance achieved at 40 mM and activity retained up to 100 mM (**Figure 6e**). This apparent discrepancy likely reflects the differing mechanistic requirements of the two assay systems. While the radiolabel incorporation assay measures nucleotide incorporation into a short, pre-activated substrate under isothermal conditions, PCR requires repeated cycles of denaturation, annealing, and extension, during which Tris may contribute to buffer capacity and stabilization of the reaction components rather than directly affecting polymerase catalysis. Yet the precise molecular mechanism underlying this phenomenon remains to be elucidated.

### 3.6 Pyrfu pol 3′→5′ exonuclease activity assay using radiolabeled nucleotides

The 3′→5′ exonuclease activity of Pyrfu pol was assessed by monitoring the degradation of [³H]-dTTP-labeled lambda/HindIII DNA over a 60 min time course, in the presence or absence of dNTPs (**Figure 5**). In the absence of dNTPs, exonuclease activity increased progressively, reaching approximately 85% at 60 min. In the presence of 100 µM dNTPs, activity was substantially reduced at all time points, reaching only approximately 40% at 60 min. This dNTP-dependent suppression of exonuclease activity is consistent with the known competitive relationship between the polymerase and exonuclease activities in family B DNA polymerases, in which dNTP binding at the polymerase active site shifts the equilibrium away from the exonuclease mode [36–37].

**Figure 5.**
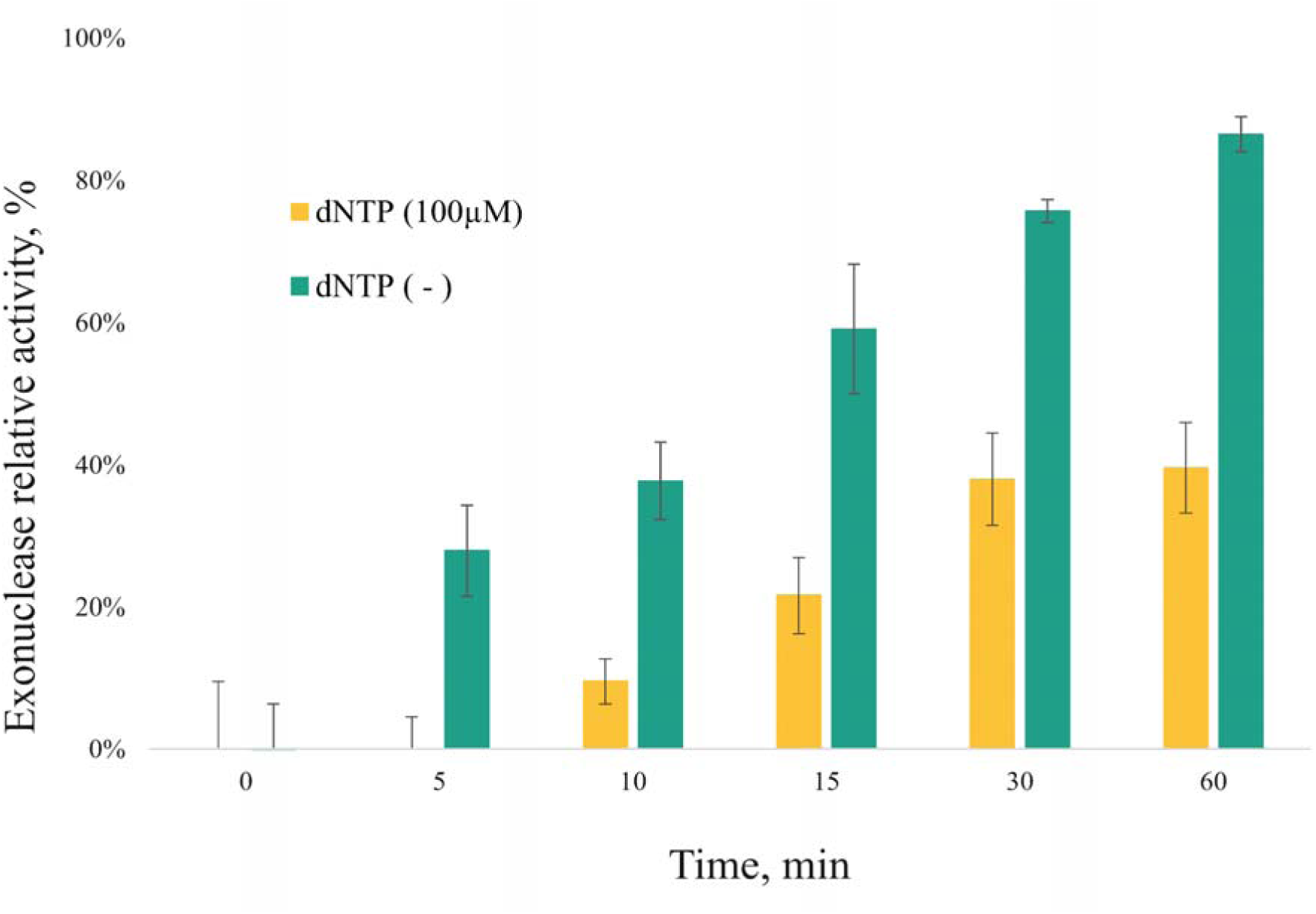
3’→5’ Exonuclease activity assay. Exonuclease activity was assessed by degradation of [³H]-dTTP lambda/HindIII DNA in the presence (gold) or absence (teal) of dNTPs over a 60 min time course.

### 3.7 Pyrfu pol activity assay using Polymerase Chain Reaction (PCR)

PCR-based optimization of Pyrfu pol reaction conditions revealed both similarities and notable differences compared with the radiolabel incorporation assay. The effect of pH on PCR performance was tested over the range of 6.0-11.0 (**Figure 6a**). A detectable amplification product was observed at pH 6.5, with optimal band intensity between pH 7.5 and 10.0, indicating remarkably broad pH tolerance under PCR conditions. This contrasts with the narrower pH optimum of 6.5-8.0 observed in the radiolabel incorporation assay, suggesting that the buffering capacity of the reaction mixture during thermal cycling may partially compensate for suboptimal pH conditions. The MgSO_4_ optimum in PCR (**Figure 6b**) was consistent with the radiolabel assay: no amplification product was detected at 0 mM, and optimal performance was observed at 2 mM, confirming the essential, concentration-sensitive role of Mg²[ ions as a cofactor [34, 38]. Activity declined gradually at concentrations above 2.5 mM, with no activity at 10 mM. The effect of KCl on PCR (**Figure 6c**) showed that amplification was absent at 0 mM, with optimal band intensity between 20 and 40 mM, and a gradual decrease at higher concentrations, with no detectable product above 80 mM. This contrasts with the radiolabel assay, in which Pyrfu pol retained over 80% of its activity across the entire 0-100 mM range, suggesting that KCl plays an additional structural role in PCR beyond its effect on polymerase catalysis alone [38]. Similarly, (NH_4_)_2_SO_4_ (**Figure 6d**) supported PCR amplification across a broad range, with optimal performance at 0-40 mM and a progressive decline at higher concentrations, with amplification lost above 50 mM. This is broadly consistent with the radiolabel assay results, where activity was retained up to 100 mM. However, the narrower PCR optimum again reflects the more stringent requirements of multi-cycle amplification. As discussed above, Tris (pH 7.0) was inhibitory in the radiolabel assay (**Figure 4f**) but essential for PCR (pH 8.5, **Figure 6e**). A minimum of 10 mM was required for barely detectable amplification, with optimal performance at 30-90 mM and activity retained up to 100 mM. Optimization of the extension rate (**Figure 6f**) showed that for short products (500 bp), all tested extension rates (15-60 s/kb, corresponding to actual extension times of 8 s, 15 s, 23 s, and 30 s) yielded comparable band intensities, indicating that Pyrfu pol can efficiently synthesize at high elongation speeds for short templates. For the 1,000 bp product, optimal amplification was achieved at 30-60 s/kb, while successful amplification of the ∼8,000 bp product required an extension rate of 60 s/kb, as indicated by the red arrow. These results establish extension rate guidelines for Pyrfu pol: 20 s for products up to 500 bp, 40 s/kb for products up to 5,000 bp, and 60 s/kb and above for longer products.

**Figure 6.**
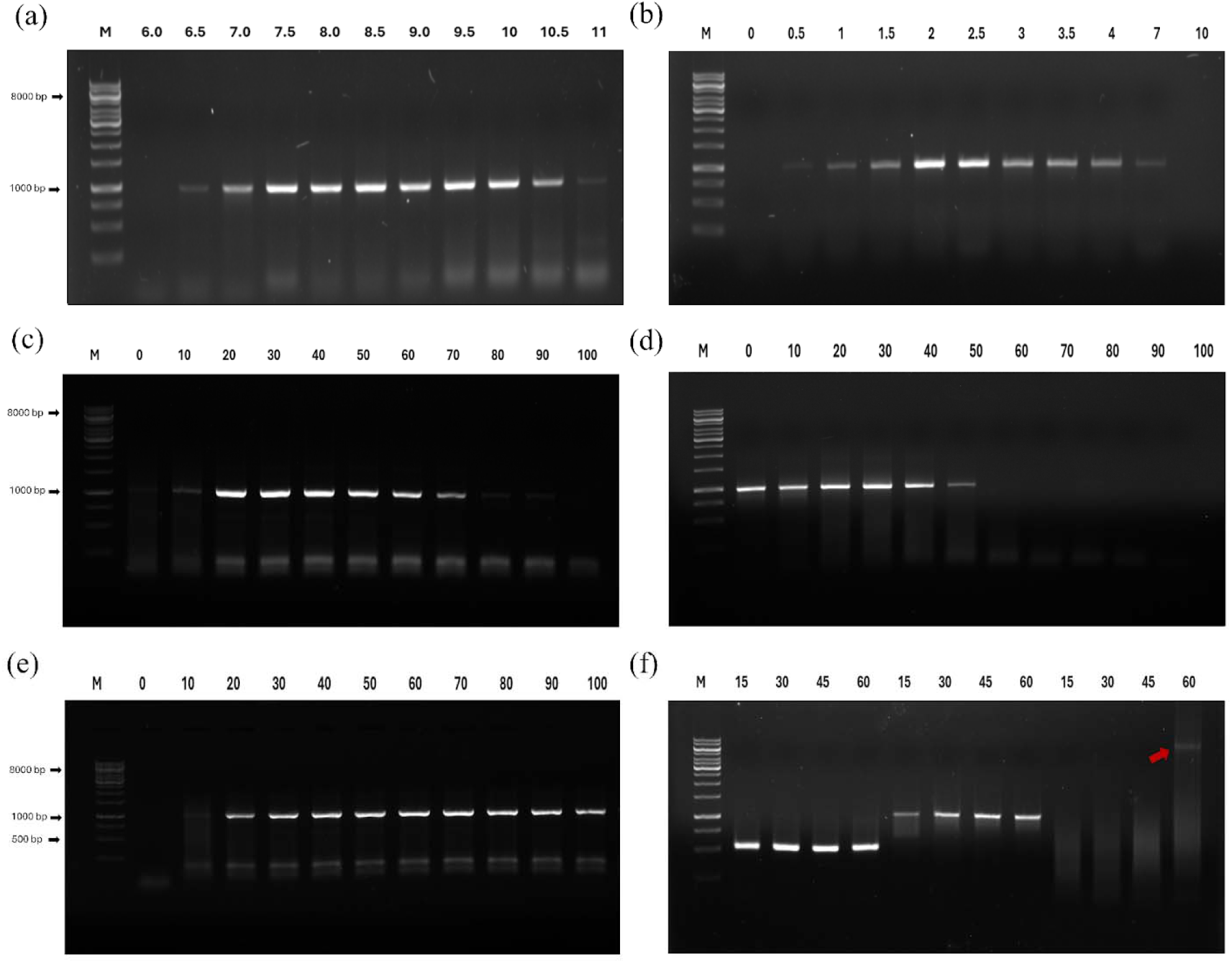
Optimization of reaction conditions for Pyrfu pol assessed by PCR. Amplification products were separated on a 0.8% agarose gel in TBE buffer and visualized by ethidium bromide staining. M: 1 kb GeneRuler DNA Ladder (Thermo Fisher Scientific); lane 1 (0 mM or lowest tested value). 500, 1,000 and ∼8,000 bp fragments were amplified using appropriate plasmid DNA as a template. Each parameter was independently varied: (a) pH (7.5-9.5), (b) MgSO_4_ (0-10 mM), (c) KCl (0-100 mM), (d) (NH_4_)_2_SO_4_ (0-100 mM), (e) Tris (pH 8.5, 0-100 mM) and (f) extension rate.

### 3.8 Analysis of Pyrfu pol processivity using PCR amplification

The processivity of Pyrfu pol was evaluated by its ability to amplify DNA fragments from 350 to 8,000 bp using pET24a(+)_Pyrfu and pET24a(+)_Pya plasmid DNA as a template (**Figure 6**). Under optimized reaction conditions, the enzyme produced detectable bands for all tested fragments up to 8,000 bp. PCR products were clearly resolved on a 0.8% agarose gel, confirming high specificity and the absence of non-specific products. This capacity for long-range amplification indicates that Pyrfu pol possesses high intrinsic processivity, a feature critical for the efficient replication of archaeal genomes under extreme conditions. These findings are consistent with the properties of other highly processive family B DNA polymerases from hyperthermophilic archaea, such as those from the *Thermococcus*, *Pyrobaculum* and *Pyrococcus* genera, which (especially the latter) are widely utilized in molecular biology for their superior performance in synthesizing long DNA templates [32,39–40].

### 3.9 Resistance of Pyrfu pol to PCR-inhibiting agents

The tolerance of Pyrfu pol to common PCR inhibitors was evaluated by amplifying a 500 bp DNA fragment in the presence of increasing concentrations of eight inhibitory compounds (**Figure 8**). Overall, Pyrfu pol showed broad tolerance to inhibitors, comparable to or exceeding that of previously characterized thermophilic family B DNA polymerases.

**Figure 7.**
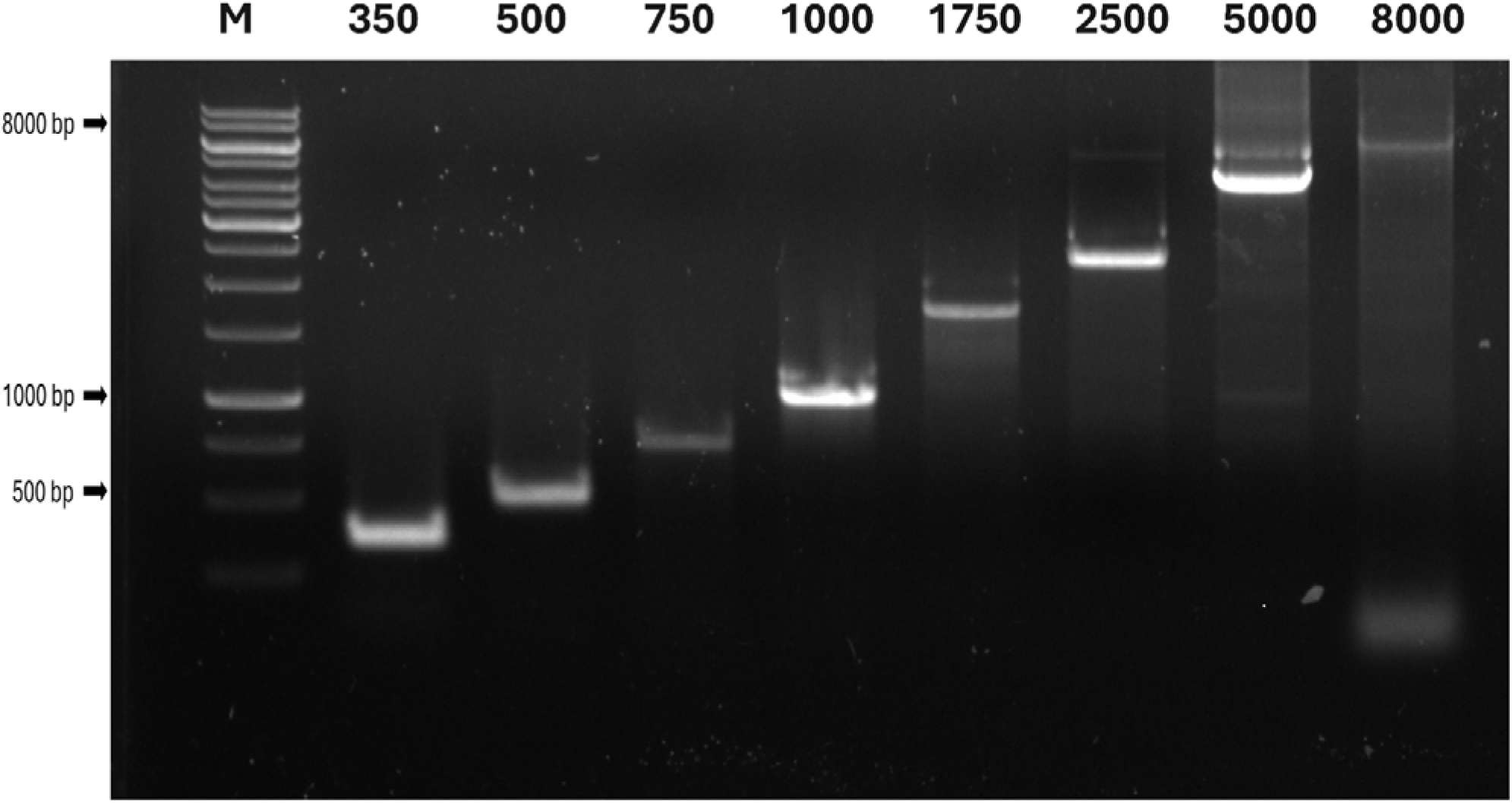
Processivity of Pyrfu pol assessed by PCR amplification of DNA fragments of increasing length. PCR products were separated on a 0.8% agarose gel in TBE buffer and visualized by ethidium bromide staining. M: 1 kb GeneRuler DNA Ladder (Thermo Fisher Scientific). Plasmid DNA pET24a(+)_Pyrfu and pET24a(+)_Pya was used as the template. Amplified fragments: 350, 500, 750, 1,000, 1,750, 2,500, 5,000, and ∼8,000 bp.

**Figure 8.**
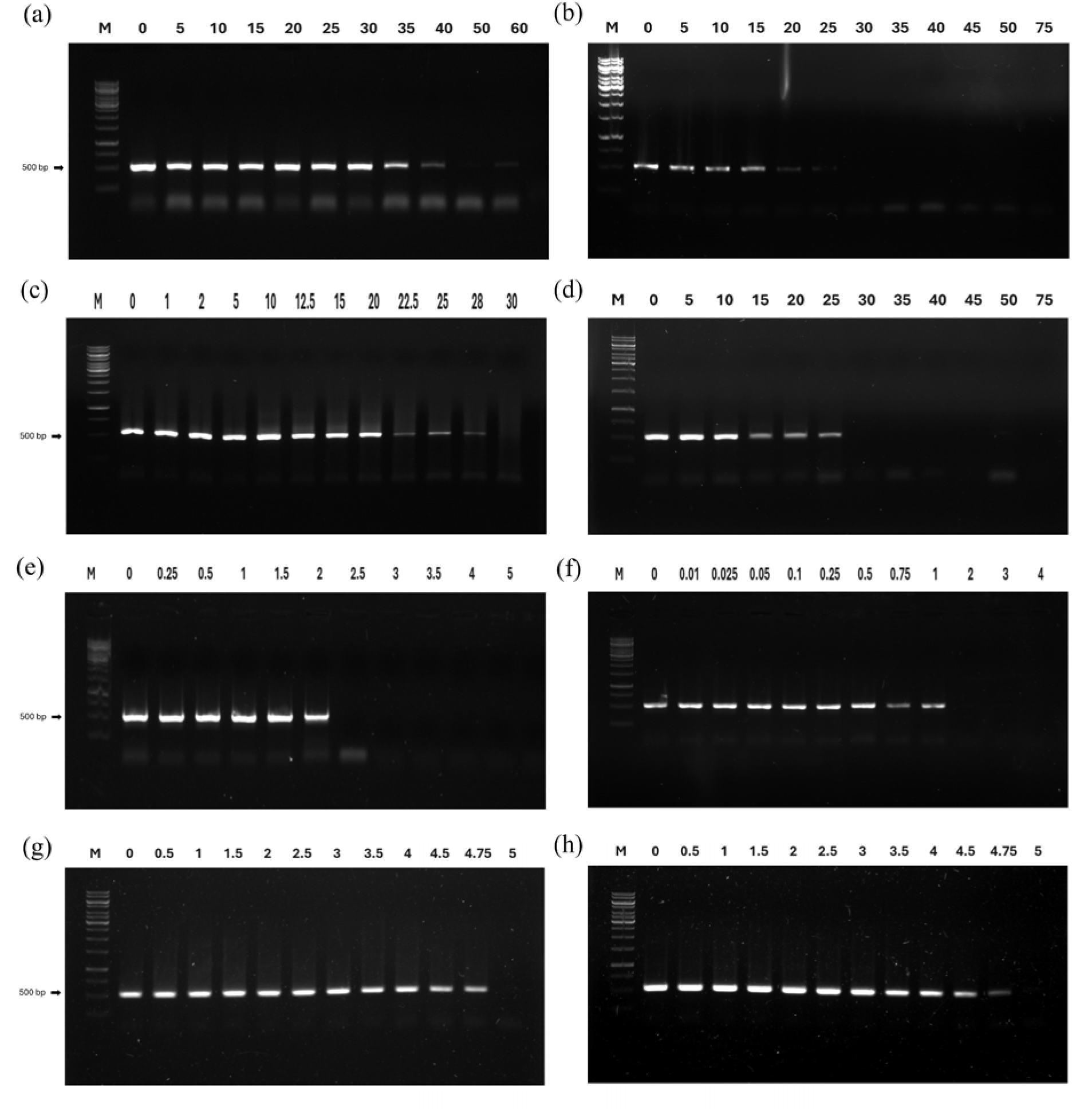
Tolerance of Pyrfu DNA polymerase to common PCR inhibitors. PCR products were separated on a 0.8% agarose gel in TBE buffer and visualized by ethidium bromide staining. M: 1 kb GeneRuler DNA Ladder (Thermo Fisher Scientific); lane 1 (0 concentration) served as a positive control. Amplification was performed in the presence of increasing concentrations of: (a) NaCl (0-60 mM), (b) heparin (0-75 ng/reaction), (c) lactoferrin (0-30 µg/reaction), (d) humic acid (0-75 ng/reaction), (e) EDTA (0-5 mM), (f) phenol (0-4% [v/v]), (g) ethanol (0-5% [v/v]), and (h) isopropanol (0-5% [v/v]). A 500 bp fragment was amplified using plasmid pET24a(+)_Pyrfu DNA as the template.

NaCl was tolerated up to 35-40 mM, with barely detectable product at 60 mM (**Figure 8a**). This tolerance is comparable to that reported for Pfu DNA polymerase, which retains activity up to 20 mM NaCl [41]. Heparin, a potent polyanion inhibitor that competes with DNA for polymerase binding, was tolerated up to 15 ng per reaction, with marginal activity at 25 ng per reaction (**Figure 8b**). By comparison, Pfu polymerase tolerates up to 125 ng/reaction [41], suggesting that Pyrfu pol is more sensitive to heparin inhibition. Lactoferrin, a glycoprotein that inhibits PCR through protein-protein interactions with the polymerase, was tolerated up to 20 µg per reaction, with marginal amplification at 28 µg per reaction (**Figure 9c**). This is comparable to the tolerance of the PfuDBD-TaqS chimeric polymerase, which is reported as 27 µg per reaction [42]. Humic acid, a common environmental inhibitor relevant to forensic and environmental DNA applications, was tolerated up to 25 ng per reaction (**Figure 8d**). EDTA tolerance was observed up to 2 mM (**Figure 8e**). This is consistent with its mechanism of action as an Mg²[ chelator.. Phenol tolerance was limited to 1% (v/v) (**Figure 8f**). Pyrfu pol also displayed tolerance to ethanol and isopropanol, retaining clear amplification up to 4.75% (v/v) for both alcohols (**Figure 8g-h**), suggesting that the enzyme maintains structural integrity in the presence of short-chain alcohols at concentrations relevant to crude DNA extract applications.

### 3.10 Polymerase fidelity assay using blue/white colony screening

The effect of the intrinsic 3′→5′ exonuclease proofreading activity on the fidelity of Pyrfu DNA polymerase was evaluated using a plasmid-based lacZα gene complementation assay [25]. As summarized in Table 2, Pyrfu DNA polymerase exhibited a mutation frequency of 2.32 × 10^-3^, corresponding to an error rate of 1.59 × 10^-5^ errors per base incorporated. In control experiments, the error rates for Taq and Pfu DNA polymerases were 4.27 × 10^-5^ and 3.91 × 10^-6^, respectively. Pyrfu DNA polymerase exhibited an error rate intermediate between the two controls, approximately 2.9-fold lower than the non-proofreading Taq DNA polymerase and approximately 5-fold higher than the proofreading-competent Pfu DNA polymerase. Although this places Pyrfu well above Taq in fidelity, its improvement over Taq is modest compared to other archaeal B-family polymerases characterized using the pJR2-lacZ assay system. For instance, Tba5 polymerase showed only a 1.39-fold improvement over Taq, comparable to Pfu in the same study (1.04-fold over Taq) [34]. In contrast, Tce, Tma, and Twa polymerases demonstrated 3.7-, 3.85-, and 3.6-fold lower error rates than Taq, respectively, again paralleling the fold-improvements observed for their corresponding Pfu controls in each study, which scored 4-, 4.12-, and 2.7-fold over Taq, respectively [37,39–40]. Notably, the Pfu fold-over-Taq values reported across these studies range from approximately 1.04- to 4.12-fold, considerably narrower than the 14.5-fold difference observed in our assay, indicating substantial inter-laboratory and inter-assay variability even for the same reference enzyme. When normalized to the Pfu control within each study, Pyrfu’s fidelity (2.9-fold over Taq, corresponding to a Pyrfu/Pfu ratio of ∼0.21, i.e., a 5-fold higher error rate than Pfu) appears modest relative to enzymes such as Tce, Tma, and Twa, whose fold-over-Taq values closely tracked or exceeded their respective Pfu controls. This suggests that, while Pyrfu retains a functional 3′→5′ proofreading exonuclease domain that confers a clear fidelity advantage over non-proofreading Taq polymerase, its overall replication accuracy remains below that of the most highly accurate family-B archaeal polymerases.

**Table 2.** Fidelities observed for DNA polymerases assessed through a plasmid-based *lacZα* gene complementation assay.

| Polymerase | Total Number of Colonies <sup>1</sup> | Number of White Colonies | Mutation Frequency <sup>2</sup> | Error Rate <sup>3</sup> | Fold improvement over Taq |
| --- | --- | --- | --- | --- | --- |
| **Pyrfu DNA polymerase** | 20,394 | **60** | $2.32 \times 10^{-3}$ | $1.59 \times 10^{-5}$ | **2.9** |
| **Taq DNA polymerase** | 19,056 | **141** | $6.77 \times 10^{-3}$ | $4.65 \times 10^{-5}$ | **1** |
| **Pfu DNA polymerase** | 22,016 | **24** | $4.65 \times 10^{-4}$ | $3.19 \times 10^{-6}$ | **14.5** |

<sup>1</sup> Colony counts represent the cumulative results obtained from four independent experiments.
<sup>2</sup> Mutation frequency was calculated as: (number of white colonies/total number of colonies) – background mutation frequency. A background mutation frequency of $6.25 \times 10^{-6}$ , determined for gapped pSJ3 plasmid DNA, was used for all calculations.
<sup>3</sup> Error rates, expressed as the number of errors per base incorporated, were derived from mutation frequencies according to previously described methods x1

### 3.11 Melting temperature analysis of Pyrfu pol by urea-nanoDSF

The thermostability of Pyrfu pol was assessed using urea gradient-nanoDSF method [26], a chemically assisted denaturation approach that enables reliable determination of melting temperatures for extremely stable proteins, as standard nanoDSF measurements failed to detect unfolding transitions. Melting temperatures were determined across a urea concentration gradient of 1-6 M and plotted as a linear function of denaturant concentration (**Figure 9**). The resulting regression line (y = -2.624x + 105.91) showed excellent linearity (R² =0.9944), confirming cooperative unfolding behavior consistent with a well-folded, structurally and kinetically stable protein. Extrapolation to 0 M urea yielded a T_m_ of 105.9 ± 0.08°C, placing Pyrfu pol among the most thermostable DNA polymerases characterized to date, consistent with the extreme thermophily of its host organism *Pyrolobus fumarii*, which grows optimally at 106 °C and tolerates temperatures up to 113 °C [16].

**Figure 9.**
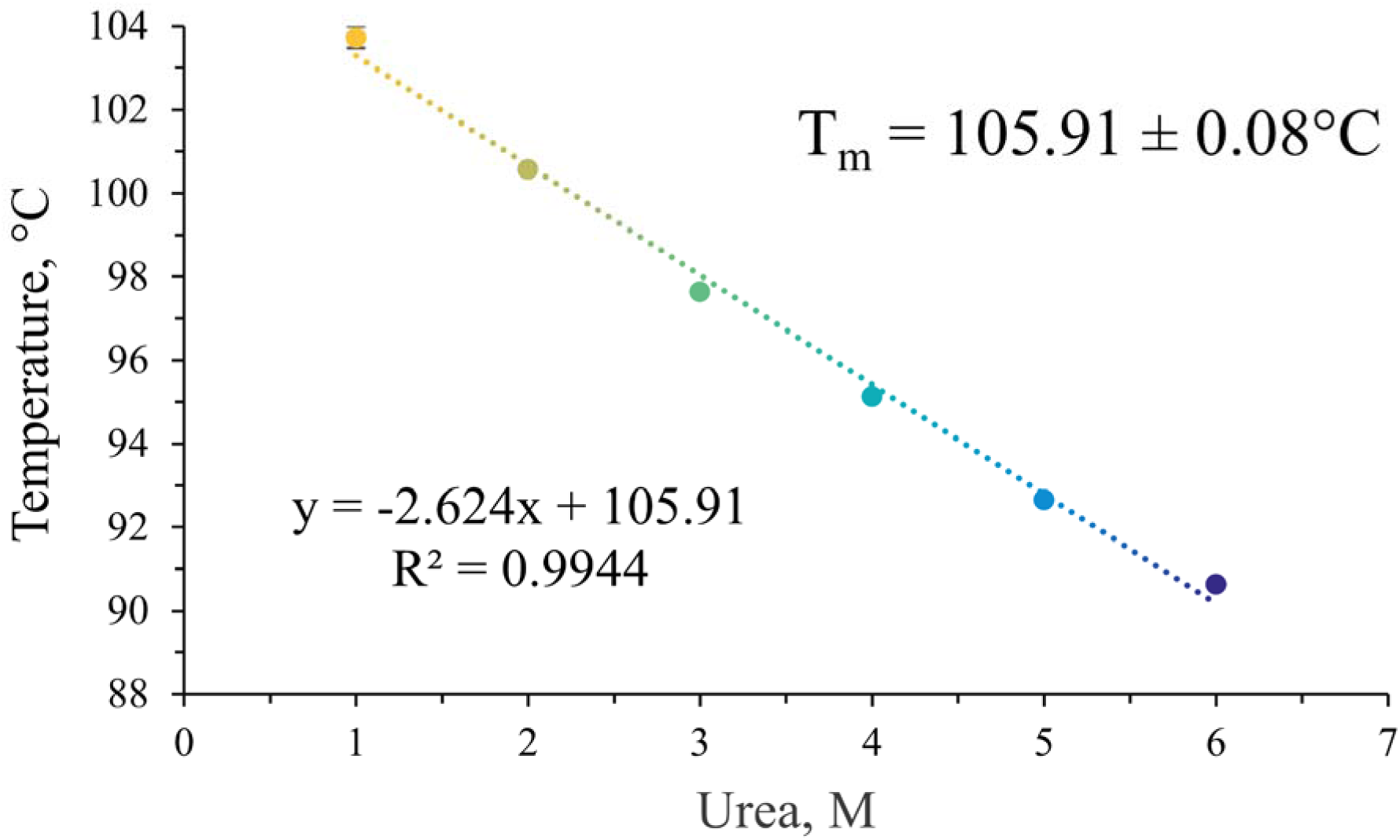
Melting temperature analysis of Pyrfu pol assessed by urea-gradient nanoDSF. The chemically assisted protein denaturation profile was presented as a linear function of polymerase T_m_ versus urea concentration (1-6 M). Each data point represents the mean T_m_ value (n=3) determined from the peak maxima of the first derivative of the F_350_/F_330_ fluorescence ratio. The intrinsic thermal stability in the absence of a denaturant was determined by linear extrapolation of the regression line to 0 M urea, yielding a T_m_ of 105.91 ±0.08 °C.

## Conclusions

In this work, we have successfully cloned, expressed, and biochemically characterized a family B DNA polymerase from the hyperthermophilic archaeon *Pyrolobus fumarii* A1, one of the most thermoresistant organisms known, with a growth optimum of 106 °C and an upper temperature limit of 113 °C. The recombinant enzyme was efficiently overproduced in *E. coli* Rosetta 2(DE3)pLysS and purified to homogeneity using a two-step procedure combining heat treatment and metal affinity chromatography. Pyrfu pol demonstrated exceptional thermostability, with a T_m_ of 105.9 ± 0.08 °C as determined by urea-nanoDSF, and retained polymerase activity across a broad range of ion concentrations, temperatures, and pH values. The enzyme exhibited an absolute requirement for Mg²[ ions and possessed associated 3’→5’ exonuclease proofreading activity, exhibiting 2.9-fold fidelity improvement over Taq polymerase. PCR optimization revealed a unique inhibitory effect of Tris in the radiolabel incorporation assay (pH 7.0), in contrast to its requirement for PCR amplification (pH 8.5). This observation has not been previously reported for archaeal family B polymerases. Pyrfu pol successfully amplified DNA fragments ranging from 350 bp to ∼8,000 bp and demonstrated tolerance to common biological and laboratory PCR inhibitors including humic acid, lactoferrin, ethanol, and isopropanol, thus highlighting its potential utility in molecular biology, forensic and diagnostic applications involving crude or inhibitor-containing DNA samples. Bioinformatic analysis, including physicochemical characterization, domain annotation, molecular prediction, multiple sequence alignment, and structural comparison with RB69 gp43, provided insight into protein structure and the molecular basis of the enzyme’s thermoadaptation. To our knowledge, this is among the first reports describing the cloning, expression, and comprehensive bioinformatic and biochemical characterization of a protein from *Pyrolobus fumarii* A1.

## Author contribution

**Wojciech Rusinek:** Conceptualization, Data curation, Formal analysis, Funding acquisition, Investigation, Methodology, Validation, Visualization, Writing - original draft.

**Sebastian Dorawa:** Formal analysis, Funding acquisition, Investigation, Methodology, Writing - original draft.

**Tadeusz Kaczorowski:** Conceptualization, Funding acquisition, Methodology, Validation, Supervision, Writing - original draft, Writing - review and editing.

## Acknowledgments

The authors used the Parula colormap for all applicable figures in this work to maintain visual consistency and enhance accessibility for readers with color vision deficiencies.

The authors would like to thank Dr. Magdalena Płotka for granting access to the Collection of Plasmids and Microorganisms infrastructure, namely the Prometheus NT.48. The acquisition of this instrument was supported by the Polish Ministry of Science and Higher Education under the Capital Equipment Grant No. 6830/IA/SP/2018 awarded to Dr. Anna-Karina Kaczorowska (University of Gdańsk, Collection of Plasmids and Microorganisms). The authors are also grateful to Dr. Aleksandra Kocot for valuable comments on the manuscript, to Kordian Zarzecki for his help with the analysis, and to Prof. Dr. Ruth A. Schmitz-Streit and her research group (Christian-Albrechts-University of Kiel, Institute of General Microbiology) for all their support and inspiring scientific experience.

## Funding

This work was supported by the University of Gdansk for financing through Research Projects of Young Scientists of the Faculty of Biology 2025 (grant no. 539-D100_B233_25) and National Science Centre, Poland, for financing under the GREIG 1 (grant no. 2019/34/H/NZ2/00584) and MINIATURA 9 project (grant no. 2025/09/X/NZ1/00075).

## Conflict of interest

The authors declare that they have no known competing financial interests or personal relationships that could have appeared to influence the work reported in this paper.

## Data accessibility

The raw and processed data that support the findings of this study are available on request from the corresponding authors.

## Notes

### Competing Interest Statement

The authors have declared no competing interest.

